# Absence of surface IgD does not impair naive B cell homeostasis and memory B cell formation in humans

**DOI:** 10.1101/332361

**Authors:** J. Nechvatalova, S.J.W. Bartol, Z. Chovancova, L. Boon, M. Vlkova, M.C. van Zelm

## Abstract

**One Sentence Summary:** Human B cells with a genetic defect in *IGHD* develop normally *in vivo*, and do not have a competitive disadvantage to IgD-expressing B cells for developing
into memory B cells.

**Abstract:** Surface immunoglobulin D (IgD) is co-expressed with IgM on naive mature B cells. Still, the role of surface IgD remains enigmatic even 50 years after its initial discovery. We here examined the *in vivo* role of surface IgD in human B-cell homeostasis and antibody responses in four individuals with heterozygous nonsense mutations in *IGHD*. All *IGHD* heterozygous individuals had normal numbers of B cells and serum immunoglobulins, and did not show signs of immunodeficiency or immune dysregulation. IgD^+^ and IgD– naive mature B cells were present in equal numbers and showed similar immunophenotypes, except for decreased expression of CD79b in the IgD– subset. Furthermore, both IgD+ and IgD– naive mature B cells had normal replication histories, similar capacities to differentiate into plasma cells upon *in vitro* stimulation, and Ig switched memory B cells showed similar levels of somatic hypermutations. Thus human B cells lacking IgD expression develop normally and generate immunological memory *in vivo*, suggesting that surface IgD might function more restricted in regulating of B-cell activation to specific antigenic structures.

## Introduction

Surface-expressed immunoglobulins (Ig) are the hallmark of B cells. During precursor development in bone marrow, each developing B cell creates an Ig molecule with unique specificity through genomic reassembly of genetic elements in their Ig loci *(1)*. This results in the expression of a receptor with the IgM isotype, which is crucial for naive B-cell survival and activation of the cell to respond to a specific antigen *(2, 3)*. In addition to IgM, circulating naive B cells co-express a receptor with the IgD isotype, resulting from alternative splicing of the exons encoding the variable domain to Cδ constant regions encoding exons *(4-6)*. IgM function has been extensively studied, and even though IgD co-expression is highly conserved in jawed vertebrates *(7)*, the biological role of surface IgD remains enigmatic even 50 years after its initial discovery *(8, 9)*.

IgD was first described in 1965 by Rowe et al. as serum Ig *(10, 11)*, prior to the identification of co-expression with IgM on the surface of B cells *(12-15)*. The IgM isotype is already expressed in progenitor B cells, either with the surrogate light chains as pre-B-cell receptor (pre-BCR) on pre-BII cells, or together with a rearranged Ig light chain on immature B cells as BCR. IgM is first expressed as pre-BCR (μ chain with surrogate light chains) by pre-B cells. IgD expression is first up-regulated after migration to the periphery at the transitional B-cell stage, and splicing to IgD is critically dependent on Zinc-finger protein ZFP318 *(16, 17)*. In mature naive B cells, the levels of IgD exceed that of IgM, and are down-regulated after antigen recognition *(18, 19)*.

Surface IgD has been implicated in regulation of tolerance induction versus antigen responses. Activation of IgD– immature B cells was found to result in tolerance induction or apoptosis, while similar antigen doses activated IgD+ mature B cells *(20)*. On the other hand, Ubelhart *et al*. recently showed using in vitro models that the large flexible hinge region of IgD prevents low valent antigens from triggering downstream signalling and B-cell activation *(21)*. This would suggest that the presence of IgD on mature B cells functions to inhibit responses to potential non-complexed autoantigens.

In terms of immune responses, IgM and IgD appear to function very similarly: both activate the same downstream signalling cascades and can both mediate B-cell activation, deletion, or anergy after interaction with specific antigen.*(22)* Furthermore, mouse models that are deficient for IgD or have IgM substituted with IgD have normal generation of B cells and are capable of generating responses to T-cell dependent and –independent antigens *(23-25)*. Still, affinity maturation is slightly delayed in the primary immune response, and the signal transduction differs qualitatively between IgM and IgD *(26)*. Together, these studies indicate potential roles for surface IgD in B-cell homeostasis and immune responses, but its function on B cells *in vivo* immunity remains unclear and prompted us to investigate this *in vivo*.

We here studied the *in vivo* function of surface IgD on human B cells in four individuals from one family who carried heterozygous germline nonsense mutations in the *IGHD* gene. Detailed clinical, molecular and cellular analyses of the family was performed, and included direct comparison of the B cells expressing the wildtype *IGHD* allele and those utilizing the *IGHD* mutant allele.

## Results

### Identification of a family with heterozygous nonsense mutations in IGHD

As part of the diagnostic work up, blood B cells were studied by flow cytometric immunophenotyping in a patient with nodular lymphocyte-predominant Hodgkin lymphoma (index; IGHD6). Peripheral blood B cells appeared polyclonal with a normal Igκ/Igλ ratio. However, the patient carried an abnormally large population of IgM-expressing B cells that lacked IgD expression (31.2% of B cells; Fig. 1A). These cells were phenotypically diverse with the majority being CD38^dim^CD27^−^ (naive), and smaller fractions being either CD38^hi^CD27^−^ (transitional) or CD38^dim^CD27^+^ (memory). Because about half of the CD27^−^ IgM^+^ naive B-cell compartment was IgD^−^, we hypothesized that these cells used an *IGH* allele with a germline mutation in the *IGHD*-encoding exons. To study whether this abnormal population was inborn, blood from the patient’s family members was immunophenotyped. The patient’s mother, maternal aunt and grandfather carried an abnormally large fraction of B cells that were IgM^+^IgD^−^CD27^−^, fitting with a monoallelic inheritance. Sequence analysis of the *IGHD* gene revealed that all four affected individuals carried the same heterozygous G>A mutation in exon 1 (Fig. 1BC). This heterozygous c.368G>A mutation results in mutation of a tryptophan (TGG) to a stop codon (TAG). Because the nonsense mutation (p.W123X) is already in exon 1, this allele will not give rise to a functional IgD molecule (Fig. 1D), resulting in the lack of IgD membrane expression in ∼50% of naive B cells in all affected individuals. The c.368G>A mutation was not present in the unaffected father and grandmother, nor in healthy control, and has not been reported in Ensembl (current as of December 2017).

**Fig 1.**
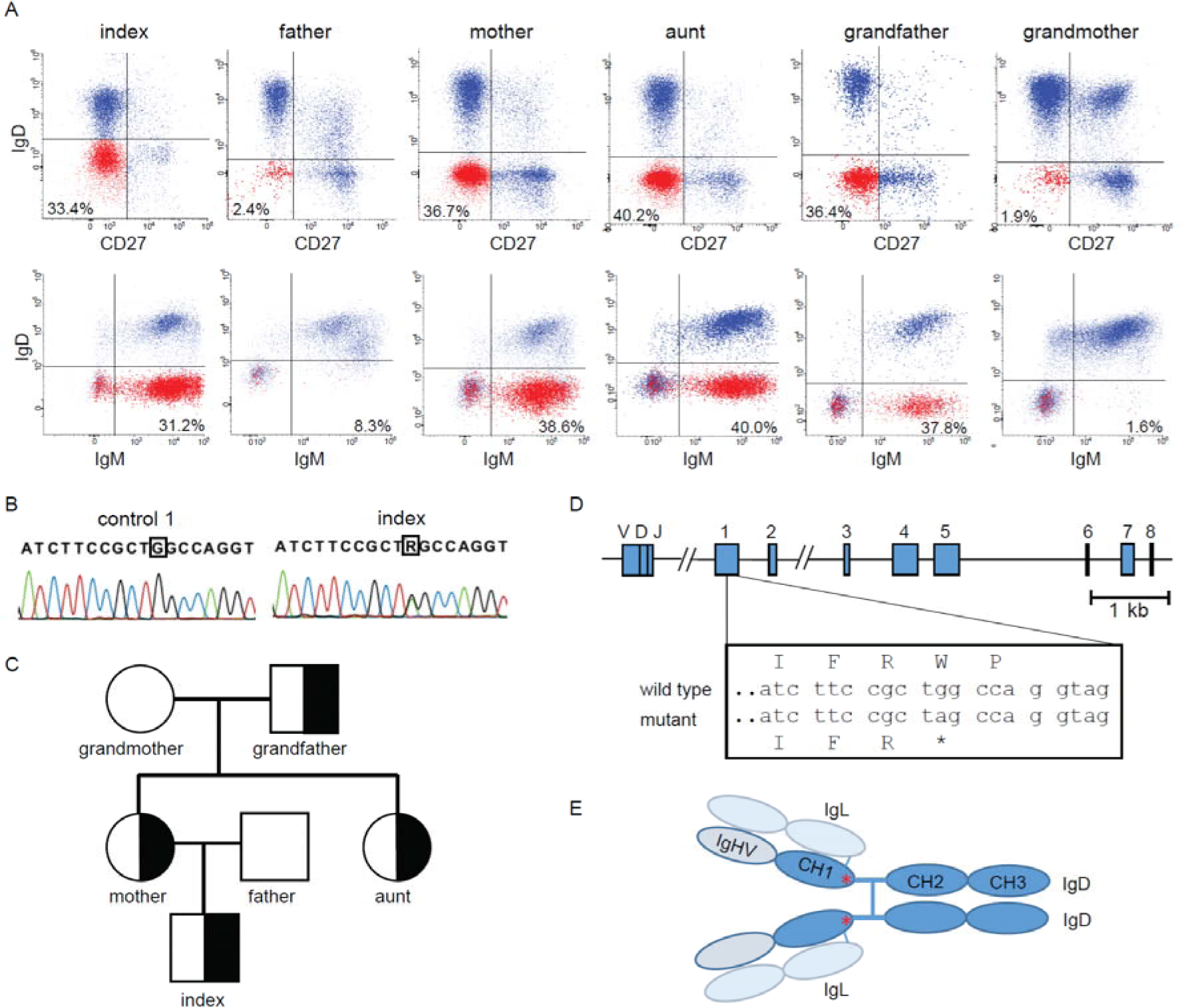
Identification of heterozygous IgD deficiency. **(A)** Large fractions of IgM-expressing B cells lack surface IgD in four individuals from one family. **(B)** Detection of a heterozygous G>A mutation in all four individuals with large IgD-negative B cell populations. **(C)** Family tree. Half-filled symbols denote known carriers of the mutation; squares denote male family members; circles denote female family members. **(D)** The c.368G>A mutation in *IGHD* exon 1 results in a premature stop codon (p.W123X) in the first Ig constant domain. IgL, Ig light chain. <u>gt</u> represents 3’ splice site of exon 1.

### Humoral immunity in IGHD heterozygous individuals

Extensive clinical and immunological work-up was performed on all 6 included family members. Except for the index case (Hodgkin lymphoma), only the affected aunt had a history of celiac disease and Turner syndrome). Besides reduced serum IgD, heterozygous IGHD-deficient individuals carried mostly normal serum Ig levels (Table 1). Specific antibody levels against previous vaccinations were normal in the two IGHD patients that were tested (Table 2). Reactivity analysis to a large panel of 13 autoantigens revealed the presence of anti-TPO autoantibodies in the grandmother, anti-GPC in the mother and anti-ENA in the father. The grandfather, aunt and index patients who are all carriers of the *IGHD* mutations were negative for all tested autoantibodies.

**Table 1.**
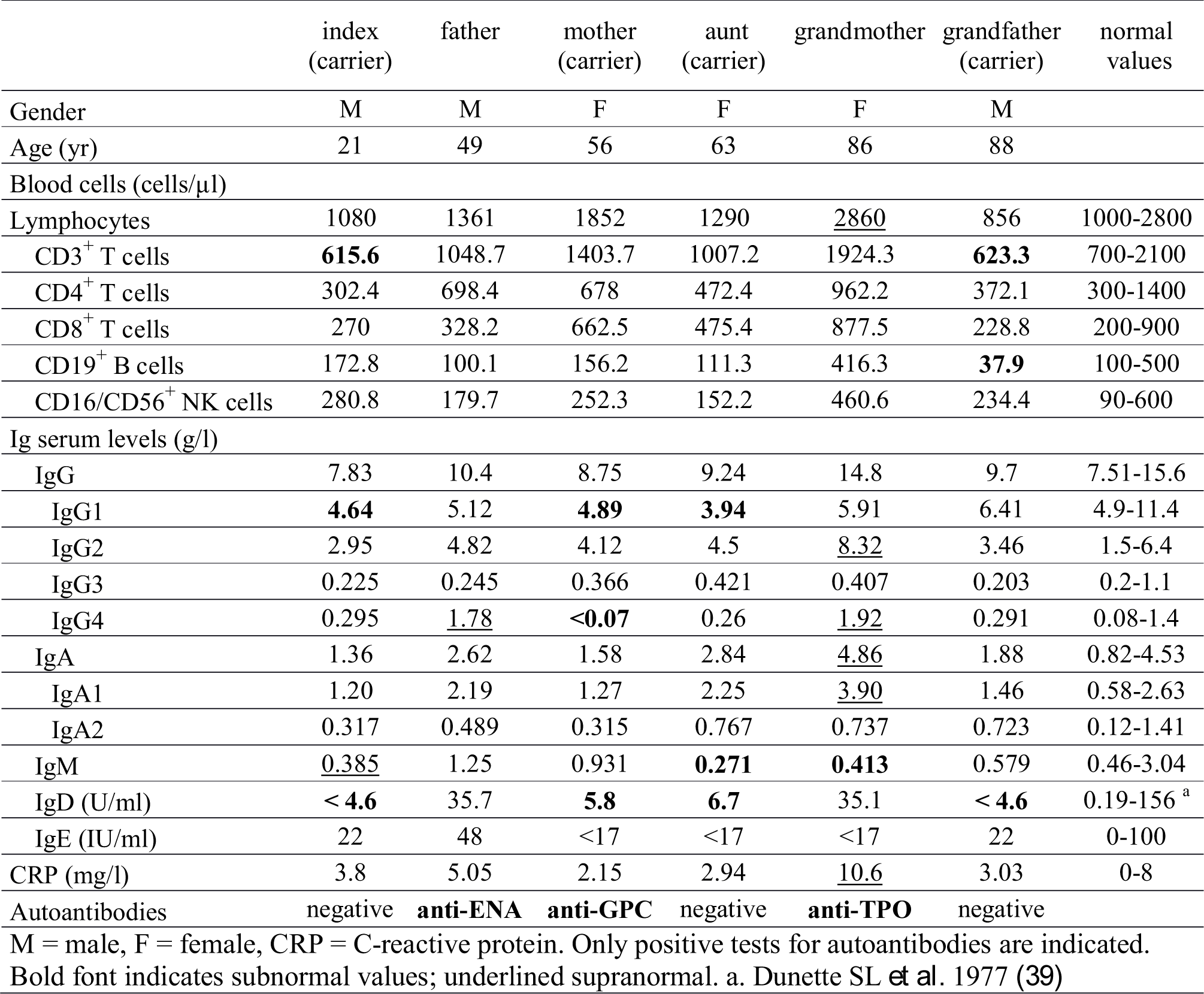
Basic and immunological characteristics of all family members

|  | index<br>(carrier) | father | mother<br>(carrier) | aunt<br>(carrier) | grandmother | grandfather<br>(carrier) | normal<br>values |
| --- | --- | --- | --- | --- | --- | --- | --- |
| Gender | M | M | F | F | F | M |  |
| Age (yr) | 21 | 49 | 56 | 63 | 86 | 88 |  |
| Blood cells (cells/μl) | | | | | | | |
| Lymphocytes | 1080 | 1361 | 1852 | 1290 | <u>2860</u> | 856 | 1000-2800 |
| CD3<sup>+</sup> T cells | **615.6** | 1048.7 | 1403.7 | 1007.2 | 1924.3 | **623.3** | 700-2100 |
| CD4<sup>+</sup> T cells | 302.4 | 698.4 | 678 | 472.4 | 962.2 | 372.1 | 300-1400 |
| CD8<sup>+</sup> T cells | 270 | 328.2 | 662.5 | 475.4 | 877.5 | 228.8 | 200-900 |
| CD19<sup>+</sup> B cells | 172.8 | 100.1 | 156.2 | 111.3 | 416.3 | **37.9** | 100-500 |
| CD16/CD56<sup>+</sup> NK cells | 280.8 | 179.7 | 252.3 | 152.2 | 460.6 | 234.4 | 90-600 |
| Ig serum levels (g/l) |  |  |  |  |  |  |  |
| IgG | 7.83 | 10.4 | 8.75 | 9.24 | 14.8 | 9.7 | 7.51-15.6 |
| IgG1 | **4.64** | 5.12 | **4.89** | **3.94** | 5.91 | 6.41 | 4.9-11.4 |
| IgG2 | 2.95 | 4.82 | 4.12 | 4.5 | <u>8.32</u> | 3.46 | 1.5-6.4 |
| IgG3 | 0.225 | 0.245 | 0.366 | 0.421 | 0.407 | 0.203 | 0.2-1.1 |
| IgG4 | 0.295 | <u>1.78</u> | **<0.07** | 0.26 | <u>1.92</u> | 0.291 | 0.08-1.4 |
| IgA | 1.36 | 2.62 | 1.58 | 2.84 | <u>4.86</u> | 1.88 | 0.82-4.53 |
| IgA1 | 1.20 | 2.19 | 1.27 | 2.25 | <u>3.90</u> | 1.46 | 0.58-2.63 |
| IgA2 | 0.317 | 0.489 | 0.315 | 0.767 | 0.737 | 0.723 | 0.12-1.41 |
| IgM | <u>0.385</u> | 1.25 | 0.931 | **0.271** | **0.413** | 0.579 | 0.46-3.04 |
| IgD (U/ml) | **< 4.6** | 35.7 | **5.8** | **6.7** | 35.1 | **< 4.6** | 0.19-156<sup>a</sup> |
| IgE (IU/ml) | 22 | 48 | <17 | <17 | <17 | 22 | 0-100 |
| CRP (mg/l) | 3.8 | 5.05 | 2.15 | 2.94 | <u>10.6</u> | 3.03 | 0-8 |
| Autoantibodies | negative | **anti-ENA** | **anti-GPC** | negative | **anti-TPO** | negative |  |

M = male, F = female, CRP = C-reactive protein. Only positive tests for autoantibodies are indicated. Bold font indicates subnormal values; underlined supranormal. a. Dunette SL *et al.* 1977 (39)

**Table 2.**
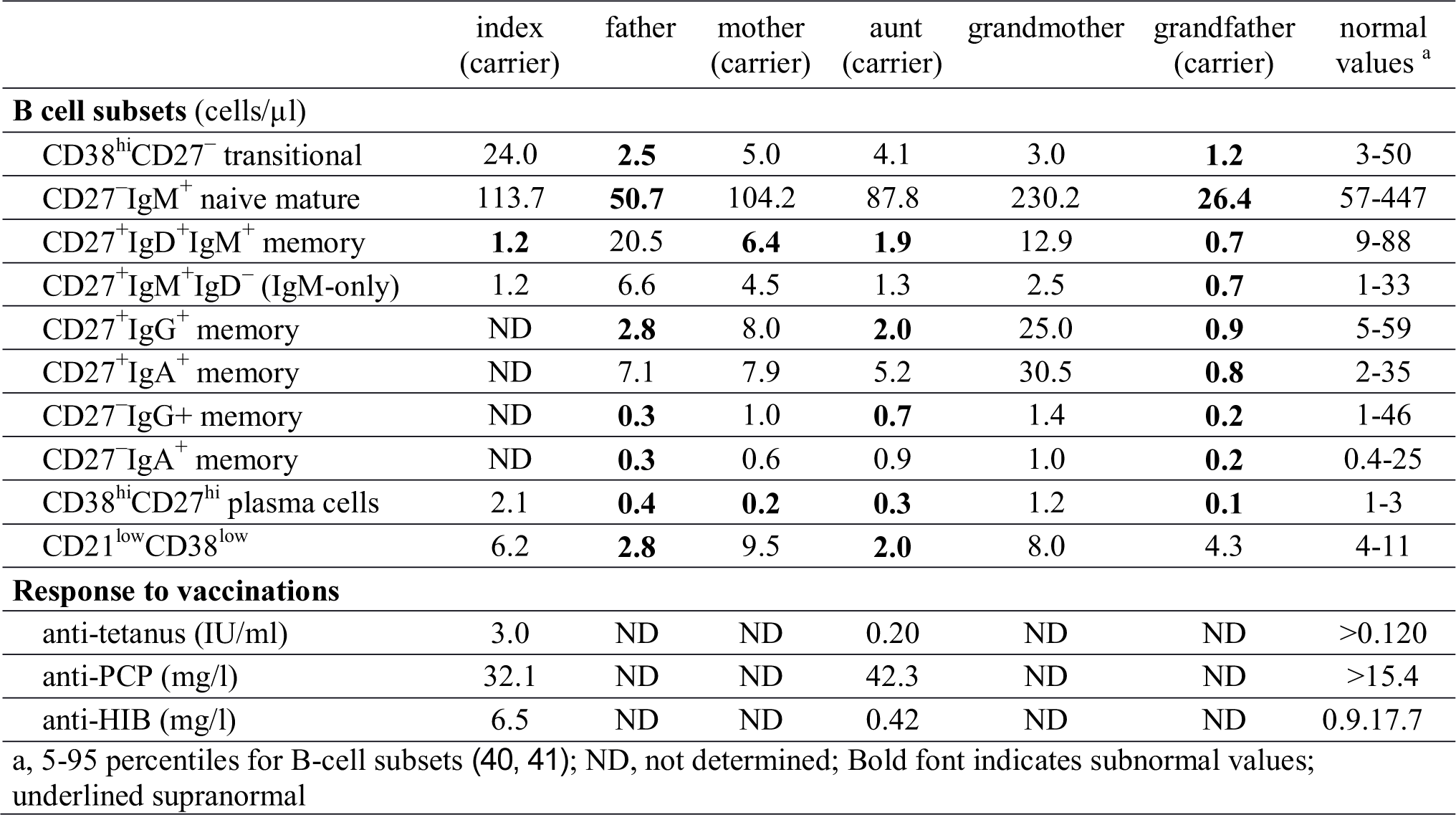
Immune response characteristics of all family members

Flow cytometric analysis revealed normal numbers of blood B cells in all individuals except for the grandfather. This reduction resulted from low numbers of both the naive and memory B-cell subsets (Table 2). Naive and memory B cells were normally present in the other individuals, except for low numbers of CD27^+^IgM^+^IgD^+^ marginal-zone like B cells in carriers of the *IGHD* mutation. To study if antigen-experienced B cells showed normal signs of molecular maturation in the *IGHD* mutation carriers, IgA and IgG transcripts were amplified from PBMC. Sequence analysis revealed the presence of multiple unique clones that nearly all contained somatic mutations. The mutation levels in IgA transcripts were slightly higher, and in IgG transcripts similar to unaffected controls (Fig. 3). All IgA and IgG subclasses were present in carriers of the IGHD mutation with slightly but significantly reduced fractions of IgA2 and IgG1 in the carriers.

**Fig 2.**
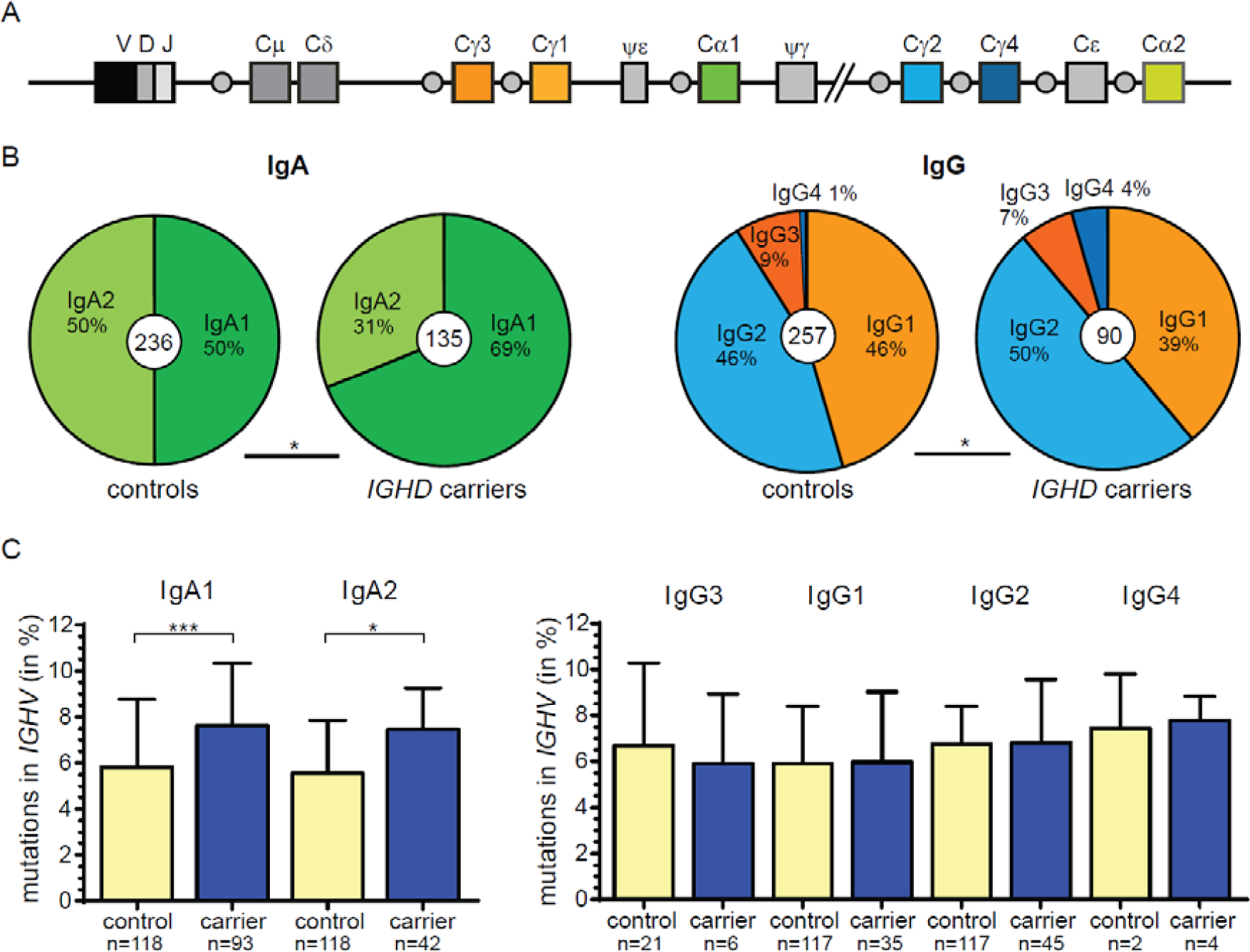
Molecular analysis of B-cell memory. **(A)** Schematic representation of the Ig constant gene regions in the human *IGH* locus. **(B)** Distribution of IgA and IgG subclass use in switched transcripts of healthy controls (n=6) and *IGHD* carriers (n=3). Total numbers of analyzed sequences are indicated in the center of each plot. Differences in the distributions were statistically analyzed with the χ2 test; *, p<0.05. **(C)** *IGHV* mutation frequencies in distinct IgA and IgG subclasses. Red lines indicate the median value. Statistical significance was calculated with the Mann-Whitney test; *, p<0.05; ***, p<0.001.

**Fig 3.**
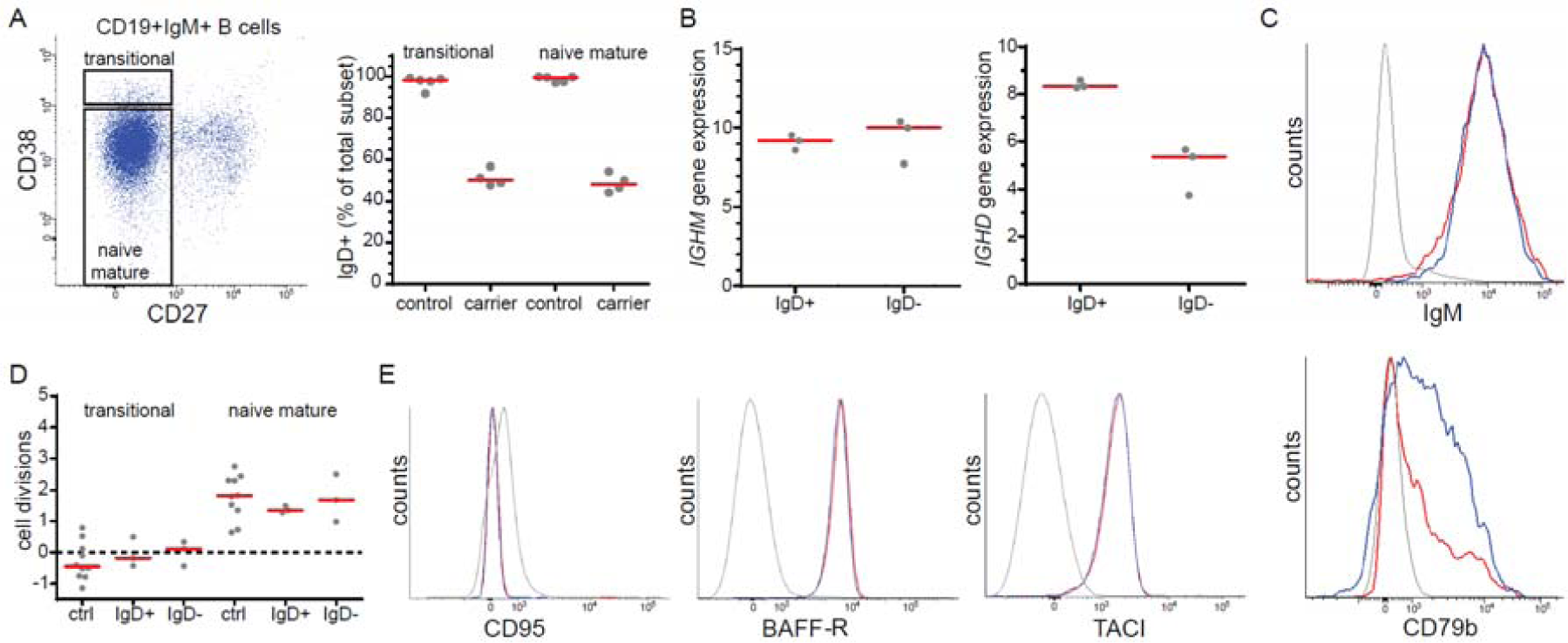
Effects of IgD deficiency on naive B cells. **(A)** Frequencies of transitional and naive mature B cells that express IgD in *IGHD* heterozygous individuals and unaffected controls. **(B)** *IGHM* and *IGHD* transcript levels in sort-purified IgD^+^ and IgD^−^ naive mature B cells from *IGHD* heterozygous individuals as determined by TaqMan-based quantitative PCR. **(C)** IgM and CD79b surface expression levels on IgD^+^ (blue lines) and IgD^−^ (red lines) naive mature B cells as determined by flow cytometry. CD3^+^ T cells were used as control (grey lines) **(D)** Replication histories of transitional and naive mature B cells from as determined with the KREC assay *(29)*. Values from control (ctrl) populations were determined previously.*(42)* Red lines represent median values. Differences between IgD- and IgD+ cells from *IGHD* heterozygous individuals and those of controls were statistically analyzed with the Mann-Whitney test. **(E)** Surface expression levels of CD95, BAFF-R, TACI on naive B cells as determined by flow cytometry (see panel C for details).

In conclusion, humoral immunity appears to be normal in carriers of the *IGHD* mutation with no overt signs of autoimmunity or immunodeficiency.

### Molecular and immunophenotypic characteristics of IgD^*–*^ naive B cells

In normal human B-cell development, IgD expression starts from the transition of immature to mature B cells, and coincides with the migration from bone marrow to the periphery *(27, 28)*. To study whether defective IgD expression affected B-cell generation or homeostasis of naive B cells, we first analyzed the frequencies of transitional and naive mature B cells expressing IgD. In unaffected controls, nearly all naive B cells in blood co-express IgM and IgD (Fig. 3A), whereas in the *IGHD* carriers this was 48.4% (Fig. 3A). These equal frequencies of IgD^+^ and IgD^−^ fractions indicate that there is no selective advantage of the presence of IgD on generation of new B cells (transitional) or homeostasis of naive mature B cells.

Naive B cells express IgM and IgD as alternative splice variants *(4)*. To study the effect of the *IGHD* nonsense mutation on IgM and IgD transcript levels, IgD+ and IgD– naive mature B cells were isolated from blood of three IGHD-heterozygous individuals. As the index patient had started treatment with rituximab, we could not obtain sufficient cells from him. IgM transcripts were equally present between subsets, whereas IgD transcripts were reduced by almost 2-fold in IgD^−^ B cells (Fig. 3B). Consequently, IgM surface expression was similar between both subsets (Fig. 3C). Importantly, surface CD79b expression levels were reduced in IgD^−^ B cells. Thus, the *IGHD* nonsense mutation does not seem to affect IgM transcript and protein levels. However, mutant *IGHD* transcript levels were reduced, and the lack of IgD surface expression resulted in a decrease in the total amount of B-cell receptors on the surface of naive mature B cells.

To further study the homeostasis of naive B cells, DNA was isolated from IgD^+^ and IgD^−^ transitional and naive mature B cells for analysis of their replication history with the KREC assay *(29)*. Similar to unaffected controls, the IgD^+^ and IgD^−^ B cells from *IGHD* heterozygous individuals did not show proliferation of transitional B cells and on average up to 2 cell divisions in naive mature B cells (Fig. 3D). Furthermore, both IgD^+^ and IgD^−^ naive mature B cells showed normal expression of BAFF-R and TACI, and lacked expression of CD95 (Fig. 3E). Thus, the absence of IgD on naive B cells does not affect their phenotype, cell numbers or replication history and consequently does not impair their generation and homeostasis.

### B-cell memory and plasma cell differentiation in absence of IgD

The individuals with heterozygous *IGHD* mutations carried Ig-class switched memory B cells. To study whether these were derived equally from IgD-expressing and IgD-deficient naive B cells, we designed a restriction enzyme assay to discriminate PCR products derived from the wildtype or the mutant allele and applied this to genomic DNA that was isolated from purified IgG and IgA memory B-cell subsets. These cells have deleted the IgD coding exons from their functionally rearranged *IGH* locus, and any amplified *IGHD* DNA would be derived from the non-functional allele. Purified IgA and IgG-switched B cells from all three carriers tested contained equal numbers of mutated and wildtype IgD alleles (Fig. 4A). Thus, *in vivo* memory B-cell formation had occurred equally from IgD^+^ and IgD^−^ naive B cells.

**Fig 4.**
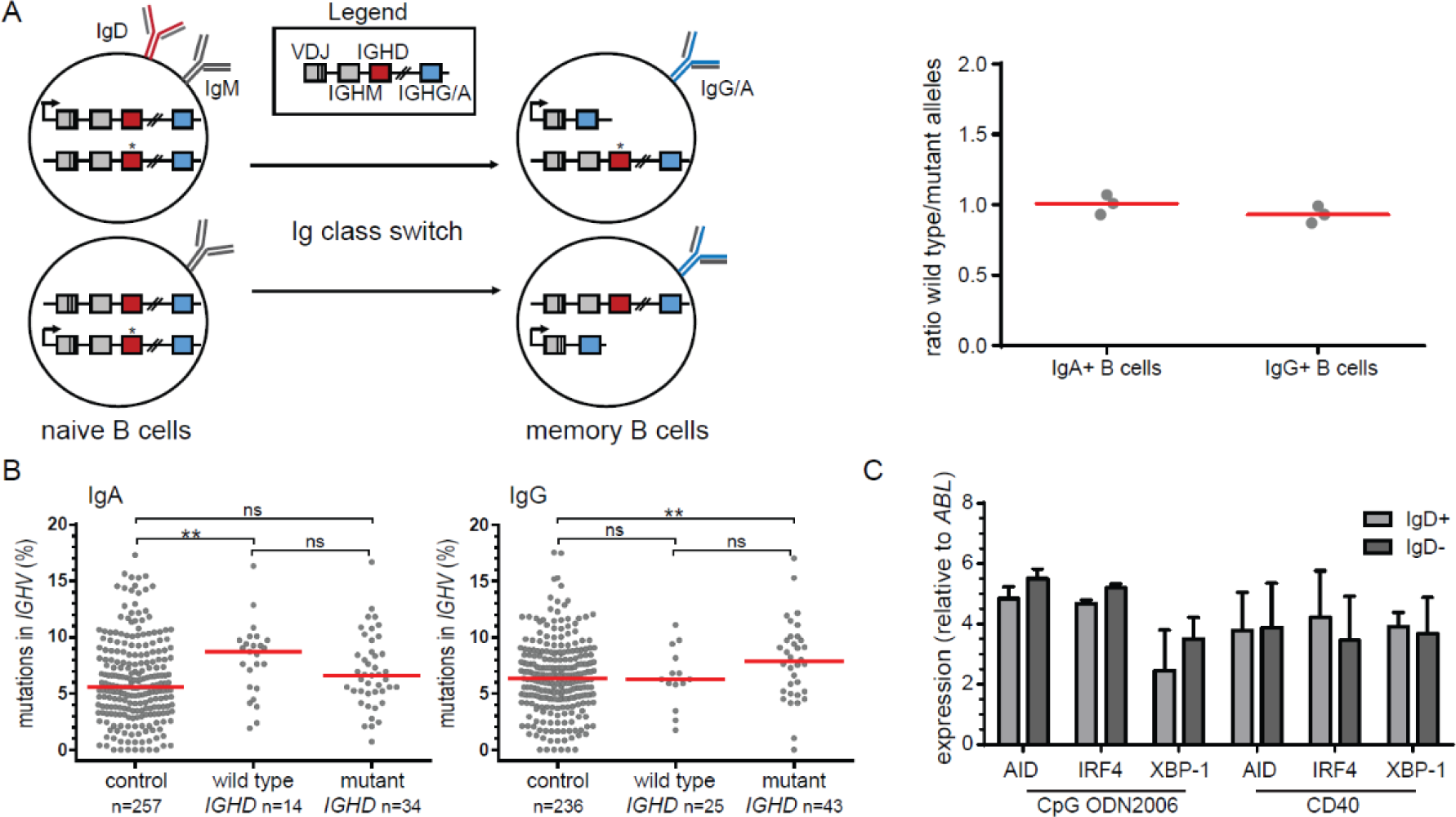
Effects of IgD on memory formation and plasma cell differentiation. **(A)** Ratio of IgA and IgG expressing memory B cells in *IGHD* heterozygous individuals that are derived from naive B cells utilizing the wild type vs mutant allele, as determined by a restriction enzyme-based assay detecting the other, unswitched allels (details in Methods). Gray dots represent ratio of wildtype and mutant (*) allele usage in IgG and IgA memory cells. **(B)** *IGHV* mutation frequencies in transcripts of Ig-class switched B cells from IgD^+^ and IgD^−^ origin. Allelic variants in *IGHV* genes were derived from from single-sorted IgD^+^ and IgD^−^ naive B-cells (**Table S4**). Red lines indicate the median value. Statistical significance was calculated with the Mann-Whitney test; *, p<0.05; ***, p<0.001. **(C)** Gene expression levels *AID, IRF4* and *XBP1* following 6 day culture of purified IgD^+^ and IgD^−^ naive B cells following stimulation with anti-IgM and CD40 or CpG. Expression was quantified relative to *ABL*. Data are expressed as the mean ± SD.

In addition to quantitative analysis of memory B-cell formation from IgD^+^ and IgD^−^ naive B cells, we performed qualitative analysis of SHM in IgA and IgG transcripts. *IGH* transcripts were sequenced from single-sorted IgD^+^ and IgD^−^ naive B cells to identify *IGHV* genes that were polymorphic between the mutant and wild type alleles of three *IGHD*-heterozygous individuals (**Table S4**). Subsequently, we designated the IgA and IgG transcripts that used these 11 alleles into those derived from *IGHD* wild type and *IGHD* mutant alleles. IgA transcripts from *IGHD* wild type alleles and IgG transcripts from *IGHD* mutant alleles carried higher SHM frequencies than those of controls (Fig. 4B).

Finally, we studied plasma cell differentiation from naive B cells in the presence or absence of IgD. Naive mature B cells from *IGHD* heterozygous individuals were sort-purified and stimulated *in vitro* with anti-IgM and anti-CD40 agonist or CpG to mimic T-cell dependent and T-cell independent secondary stimulation. After 6 days of culture, IgD^+^ and IgD^−^ B cells expressed similar levels of activation-induced cytidine deaminase (AID), as well as transcription factors IRF4 and XBP1 (Fig. 4C).

In conclusion, the IgD^+^ and IgD^−^ naive B cell in heterozygous *IGHD*-deficient individuals show equal homeostasis and are similarly capable of differentiation into memory and plasma cells.

## Discussion

In this study, we described four members from one family with heterozygous nonsense mutations in *IGHD*. Despite half of their B cells lacking IgD expression and reduced serum IgD levels, these individuals did not display overt clinical or immunological defects. Our detailed analysis showed that the lack of functional IgD did not impair B-cell generation, homeostasis or differentiation into memory B cells and plasma cells.

The expression of IgM was not affected in transitional and naive mature B cells that utilized the IgD mutant allele. As a result, these cells carried fewer surface Ig molecules than IgD-expressing B cells. In the early 1990s, two IgD-deficient mouse models were generated *(24, 25)*. The IgD-deficient B cells in these models showed increased surface IgM levels, resulting in normal total surface Ig. Although the lack of compensation by increased IgM in the human IgD-deficient B cells could reflect a difference in species, it is more likely that the difference results from the introduced mutations. IgD transcript levels in B cells from mutant mice were strongly reduced, most likely due to increased splicing of the rearranged VDJH exon to the first Cμ coding exon at the expense of splicing to Cδ *(24, 25)*. This is recapitulated in *zfp318* mutant mice of which the B cells are impaired in splicing of VDJH to *IGHD* and express threefold more surface IgM *(16)*. The *IGHD* p.W123X mutation did result in reduced IgD transcripts levels. Because this was not compensated by increased IgM transcript, this is likely a post-splicing effect of transcript instability due to the premature stop codon.

The expression of surface Ig is crucial for B-cell survival: *in vivo* ablation of surface Ig from mouse B cells was found to result in upregulation of surface Fas (CD95), which rendered these cells susceptible to T-cell mediated killing *(2)*. Although B-cell survival can be directly affected by surface Ig levels, we did not find evidence for this in IgD-deficient B cells. Transitional and naive mature B cells in the four *IGHD* heterozygous were equally composed of IgD^+^ and IgD^−^ B cells. Furthermore, CD95 expression levels were not increased, nor were the replication histories, thereby excluding compensatory proliferation. Thus, the absence of IgD and the lower levels of surface Ig did not appear to affect *in vivo* human naive B-cell homeostasis. This contrasts previous observations from mice heterozygous for the previously mentioned *IGHD*-deficient alleles *(24, 25)*, as these carried ∼2 times more B cells expressing the wild type IgD allele than B cells expressing the IgD mutant allele. On the other hand, *zfp318* null B cells that lack IgD expression have increased surface IgM levels and these are present in normal numbers in blood and spleen *(16)*. Still, population of peripheral lymphoid compartments by *zfp318*-deficient B cells has not been tested in a competitive setting with wild type B cells. Thus, it is possible that it is not the absence of IgD but the overexpression of IgM that might impair naive B-cell survival in IgD-deficient murine B cells. Irrespective, our studies demonstrate that the lack of surface IgD does not impair homeostasis of naive B cells in humans.

On top of normal naive B-cell homeostasis, we did not find evidence of impaired memory B-cell or plasma cell formation from IgD-deficient B cells either. IgA and IgG memory B cells were equally derived from *IGHD* wild type and mutant expressing naïve cells, and carried similar levels of SHM. Furthermore, IgD^+^ and IgD^−^ naive B cells were equally efficient in differentiation into plasma cells following stimulation with anti-CD40 agonist or CpG. IgD-deficient mice showed mostly normal humoral immunity as well *(24, 25)*. Still, in heterozygous mutant mice, specific IgG1 generated following immunization with a protein antigen was predominantly produced by cells expressing the IgD wild type allele. We were not able to detect IgG variants in the *IGHD* carriers, preventing the analysis of antigen-specific responses of IgD^+^ and IgD^−^ B cells. Possibly, IgD-deficient B cells respond differently to specific types of antigens, e.g. polyvalent antigens (see below) *(21)*. Still, this will be restricted to a relatively small amount of antigens, as we did not observe selective involvement of IgD wild type alleles in antigen-experienced B cells *in vivo*.

Structurally, IgD differs mostly from IgM by the presence of a large hinge region. Ubelhart *et al.* recently demonstrated that the hinge region makes IgD non-responsive to monovalent antigens *(21)*. Binding of polyvalent antigens to surface IgD normally activated the cells and did not result in an overresponse. These findings support a function of IgD to limit rather than enhance certain types of responses. This could be important in suppression of responses to autoantigens. However, we did not find evidence of enhanced autoimmunity in our patients. Following testing of 13 types of autoantibodies, only 2/4 *IGHD* heterozygous individuals were positive to one each. This suggests that surface IgD only has a limited or no function to prevent autoimmunity. However, to definitively rule this out, more individuals would need to be studied.

The lack of functional impairments in human B cells expressing a defective *IGHD* allele was not expected, because surface IgD expression is highly conserved in jawed vertebrates *(7)*. If it does not confer a general advantage for homeostasis or antigen responses of naive B cells, what would be the function of surface IgD? We were not able to study the complete absence of IgD *in vivo*, as we did not find an individual with biallelic IgD mutations. Therefore, in all our study subjects, deficits of IgD-deficient B cells to specific antigens could have been obscured by IgD-expressing B cells.

In conclusion, in the absence of surface IgD, human B cells are normally produced and capable of antigen-dependent maturation that is not outcompeted by IgD+ B cells as assessed in our assays. Rather than affecting the “wiring” of B-cell differentiation and maturation, surface IgD might have a more specific role in modulating responses to antigen based on their structural organization.

## Materials and Methods

### Patients

This family came to our attention when the index (IGHD6; Table 1) was referred to the Department of Immunology and Allergology of the St. Anne’s University Hospital in Brno on the basis of having enlarged inguinal and axillar lymph nodes. Histological examination proved granulomatous mycotic lymphadenitis. Later, he was diagnosed as nodular lymphocyte-predominant Hodgkin lymphoma (clinical stage III A). Five other members of the family (father; mother; and aunt, grandmother and grandfather from mother’s side) underwent extensive immunological examinations after informed consent was obtained and according to the guideline of the local medical ethics committee. They had no clinical manifestation of recurrent infection and/or malignancy. The aunt suffered from celiac disease and Turner syndrome.

### Flow cytometric immunophenotyping and purification of B-cells from human blood

All peripheral blood samples were obtained with informed consent and according to the guidelines of the Medical Ethics Committee of Erasmus MC and the Institutional Review Board of St. Anne’s University Hospital.

Absolute counts of blood CD3^+^ T cells, CD16+/56^+^ natural killer (NK-)cells, and CD19^+^ B cells were obtained with a diagnostic lyse-no-wash protocol. For detailed 11-color flow cytometry, red blood cells were lysed with NH_4_Cl prior to incubation of 1 million nucleated cells for 15 minutes at room temperature in a total volume of 100μL (antibodies listed in **Supplemental Table S1**). After preparation, cells were measured on a 4-laser LSRFortessa flow cytometer (BD Biosciences) using standardized settings *(30)*. Data were analyzed with FACSDiva software V8.0 (BD Biosciences).

### DNA isolation and IGHD mutation analysis

DNA was isolated from post-Ficoll granulocytes and sorted B-cell subsets with the GenElute mammalian genomic DNA miniprep kit (Sigma-Aldrich). All 8 *IGHD* encoding exons including splice sites (NCBI NG_001019) were PCR amplified (**Table S2**) from granulocyte DNA and sequenced on an ABI Prism 3130 XL fluorescence sequencer (Applied Biosystems).

### Sequence analysis of complete IGH gene rearrangements

IgD+ and IgD-naive mature B cells (CD19+IgM+CD27–CD38^dim^) were single-cell sorted into 96-well PCR plates containing 4 μl lysis solution (0.5× PBS containing 10mM DTT, 8 U RNAsin (Promega), and 0.4 U 5′-3′ RNase Inhibitor (Eppendorf)) and immediately frozen on dry ice. RNA from single cells was reverse-transcribed in the original 96-well plate in 12.5 μl reactions containing 100 U Superscript III RT (Life Technologies) for 45 minutes at 42°C using primers in the leader sequence of *IGHV* subgroups and a Cμ reverse primer *(31-33)*.

IgA and IgG transcripts were amplified from the cDNA of post-Ficoll mononuclear cells by using the same *IGHV* subgroup-specific forward primers in combination with a Cα or Cγ consensus reverse primer *(32, 34)*. The usage of V, D, J genes as well as the junctional regions were analyzed using the international ImMunoGeneTics (IMGT) information system (http://imgt.cines.fr/) *(35)*. IgG and IgA subclasses were identified by using the germline sequence of the *IGH* locus (NG_001019).

### Replication history analysis using the KREC assay

The replication history of sorted IgD+ and IgD– transitional (CD19^+^CD27^−^IgM^+^CD38^hi^) and naive mature (CD19^+^CD27^−^IgM^+^CD38^dim^) B-cell subsets was determined with the Kappa-deleting Recombination Excision Circles (KREC) assay as described previously *(29)*. Briefly, the amounts of coding and signal joints of the *IGK*-deleting rearrangement were measured by RQ-PCR in DNA from sorted B-cell populations on an ABI Prism 7000 (Applied Biosystems). Signal joints, but not coding joints are diluted two-fold with every cell division *(29)*. To measure the number of cell divisions undergone by each population, we calculated the ratio between the number of coding joints and signal joints. The previously established control cell line U698 DB01 (InVivoScribe) containing one coding and one signal joint per genome was used to correct for minor differences in efficiency of both RQ-PCR assays.

### Restriction-enzyme based assay for IGHD allele usage

Ig-class switched cells have deleted *IGHD* from the functional *IGH* allele, and only the non-functional *IGHD* allele can be amplified by PCR. Since, the nonsense mutation in exon 1 of *IGHD* disrupts an MscI restriction site (tggcca), exon 1 from IGHD was amplified with a FAM-labeled forward primer from DNA of purified CD19^+^CD38^dim^IgA^+^ and CD19^+^CD38^dim^IgG^+^ memory B cells from heterozygous IGHD-deficient individuals. The PCR products were digested with MscI (New England Biolabs) and run on the ABI Prism 3130 XL. The relative amounts of uncut (mutated) *IGHD* were compared with those in granulocytes that each carry both *IGHD* alleles. An increase in undigested product would suggest more frequent usage of the wild type *IGHD* allele in the Ig-class switched cells. DNA from healthy controls was used as a positive control for complete digestion with MscI.

### In vitro *plasma cell differentiation of purified naive B cells*

IgD^+^ and IgD^−^ naive B cells were purified and cultured with combinations of anti-IgM F(ab’)2, anti-CD40 agonist, CpG ODN2006, and IL-21, as described previously *(36, 37)*. Cells were harvested after a 6-day culture for TaqMan-based quantitative RT-PCR on a StepOnePlus (Applied Biosystems). Target gene expression levels were determined in freshly isolated and cultured cells with intron-spanning primes and fluorogenic probes (**Table E3**) and expressed relative to the *ABL* control gene *(38)*. All quantitative RT-PCR reactions were performed in duplicate.

### Serology

Using a nephelometric method, the levels were determined of CRP, IgG, IgA, IgM and IgG subclasses (Image 800 Immunochemistry System, Beckman Coulter), as well as IgD, IgE and the IgA subclasses (Behring Nephelometer II, Siemens, Marburg, Germany). Enzyme-linked immunosorbent assay (ELISA) was used for the determination of specific antibodies: anti-extra nuclear antigens (ENAscreen, BL Diagnostika GmbH, Mainz, Germany), anti-thyroid peroxidase (AESKULISA a-TPO, Aesku.Diagnostics, Wendelsheim, Germany), anti-thyroglobulin (AESKULISA a-TG, Aesku.Diagnostics), anti-tissue transglutaminase (AESKULISA tTg-G, and AESKULISA tTg-A, Aesku.Diagnostics), and rheumatoid factor (AESKULISA Rf-AGM, Aesku.Diagnostics). Indirect immunofluorescence (IIF) was used to detect anti-nuclear antibodies (HEp-2 kit, Euroimmun, Lübeck, Germany); anti-neutrophil cytoplasmatic antibodies (Granulocytes (EOH), Euroimmun) and anti-gastric parietal cells, anti-reticulin, and anti-smooth muscle antibodies (Autoantibodies RL/RK/RS kit, Orgentec, Mainz, Germany).

### Statistical analyses

Statistical analyses were performed with the Mann-Whitney U test, paired student’s T-test, or χ^2^ test as indicated in detail in Figure legends. p values <0.05 were considered statistically significant.

#### Supplementary Material

Table S1. Antibodies used for flow cytometry

Table S2. Primer sequences for PCR amplification and sequencing of IGHD coding regions and splice sites from genomic DNA

Table S3. Primers and probes used for quantification of gene transcripts

Table S4. Polymorphic *IGHV* alleles in IGHD carriers

Fig S1. Sequence detection of the c.368G>A mutation in *IGHD* exon 1.

## Acknowledgements

This work was supported by grants 15-28732A and 15-28541A of the Czech Health Research Council, the LLP/Erasmus programme, and by a National Health and Medical Research Council Senior Research Fellowship (GNT1117687) to MCvZ.

